# Distribution and Effect of *Echium Plantagineum* L. on Floral Species Diversity in the Dejen District, Northern Ethiopia

**DOI:** 10.1101/2025.06.16.660053

**Authors:** Getasew Kefale Tamiria, Kidist Belayneh Meteko

## Abstract

*Invasive plant species are considered the greatest global threat to the loss of biodiversity. Echium plantagineum* L. *an invasive species, is primarily found along roadsides and grazing lands in the Amhara Regional State of the Dejen district. The purpose of this study was to investigate the distribution and effect of E. plantagineum on the diversity of floral in the area of its occurrences. Reconnaissance survey was conducted to observe the distribution, spread, and effect of E. plantagineum on plant and animal species. As a result, 116 systematically established sample plots from the study were taken (58 invaded and non-invaded plots for each). To gather information on trees (shrubs) 10m x 20m sample plots were created on grazing land types. At each corner, a 1m^2^ subplot was taken to collect herbaceous plants. A 4m^2^ plot on roadside land units was created to gather information on the herbaceous flora, tree or shrub seedlings. Plants in each plot were recorded and named. Visual estimation was used to determine the percentage cover of herbaceous species. The study assessed species diversity using species evenness, Shannon Diversity Index, and Simpson Index of Dominance measures. These metrics were analyzed using PAST software. Similarity between invaded and non-invaded areas was evaluated using Jaccard’s similarity index and beta diversity. The relationship between the abundance of E. plantagineum and floral species abundance was examined through linear regression and correlation analyses. A total of 85 plant species from 32 families were discovered in the study areas. In non-invaded land units, 80 plant species from 30 families were recorded. In comparison, the invaded area contained 54 plant species grouped into 24 families. Thus, the number of plants was reduced by 32.5% in the E. plantagineum-invaded area compared to the non-invaded area. The findings also showed that the non-invaded areas have a higher Shannon Diversity index value was observed in plant (3.537) and also higher plant species evenness value (0.429) compared to the E. plantagineum invaded areas. The study also showed a negative relationship between the abundance of E. plantagineum invasives and the abundance of native plant species per study plot. It was concluded that E. plantagineum, the invasive plant species in the study area, has reduced floral species diversity. Therefore, it is strongly advised to implement adequate planning and strategies to detect or stop the spread and effects of E. plantagineum. This can be achieved by creating communication channels between the regional, zonal, and district agricultural offices*.

## 1. Introduction

Invasive plant species (IPSs) are non-native or alien to the ecosystem and their introduction can lead to economic or environmental harm (Byabasaija *et al*., 2020). The invasion of these species is considered as the greatest global threat to biodiversity (Amare Seifu *et al*., 2017). *Echium plantagineum* L. (Paterson’s curse or Salvation Jane) is an annual or biennial species, Native to Northwest Africa, the Iberian Peninsula, and Atlantic Western Europe; it is a noxious weed in Australia. *Echium plantagineum* L. was first introduced in Australia as an ornamental and deliberately spread in the mid-nineteenth century (Shea *et al*., 2000). It is now common, and frequently abundant throughout southwest Western Australia, central and southern South Australia, southeast Queensland, central and southern New South Wales, Victoria, Tasmania, and the Northern Territory. At the edge of its range, it is mostly limited to disturbances on roadsides. In New Zealand, *E. plantagineum* is abundant north of the volcanic plateau of the North Island and scattered to rare elsewhere (Ansari *et al*., 2016).

It produces toxic pyrrolizidine alkaloids in the shoots (Skoneczny *et al*., 2015) and bioactive shikonins in the root (Lloyd, 2011; Zhu *et al*., 2016). It competes with pasture grasses and is often toxic to grazing livestock. Globally, *E. plantagineum* is currently infesting more than 30 million hectares of land, leading to annual losses that exceed A$250 million (Zhu *et al*., 2017). Introduced species are of global concern in terms of their inherent economic and environmental costs, with annual losses of $1.4 trillion associated with biological invaders around the world (Bai, 2014; Hall, 2019). *Echium plantagineum* L. is self-incompatible in its native range but purportedly became self-compatible after its introduction to Australia (Petanidou *et al*., 2012). The plant, which is present on more than 33 million hectares of Australian land (Piggin & Sheppard, 1995), is thought to harm upwards of $30 million (AUD) annually (Shea *et al*., 2000).

Invasive plant species (IPS) have significant social, ecological, and economic impacts which cause obvious changes in artificial or natural ecosystems. They pose significant challenges to agricultural lands, range lands, biodiversity, national parks, waterways, rivers, power dams, roadsides, and urban green spaces. Their presence has both economic and ecological consequences, impacting grazing areas and facilitating the spread of vector-borne diseases (Gedyon Tamiru, 2017; Wakshum Shiferaw *et al*., 2018). From their threat to biodiversity and ecosystem services, invasive alien plant species have significant social, ecological, and economic impacts (Mohammed Mussa *et al*., 2018).

Invasions of invasive alien plants (IAPs) serve as a hiding place for crop pests, and wild animals, and contribute to the spreading of vector-borne diseases (Mohammed Mussa *et al*., 2018). *Echium plantagineum* L. is an alien invasive species, that may spread and grow vigorously when introduced deliberately or unintentionally by gardeners, traders, and foresters (Singh *et al*., 2011) and threaten biodiversity, ecosystems, economy, and human health (Levine *et al*., 2003).

*Echium plantagineum* L. is an annual herbaceous species, that belongs to the Boraginaceae family, it grows from 30 to 80cm in height, has rosette leaves that are up to 25cm long, and has very showy purple flowers that grow about 2 to 3cm in length arranged in an erect raceme (Weber, 2017). Africa may be principally susceptible to exotic and invasive species colonization due to its climate-sensitive distribution of native flora and fauna. In Ethiopia, IAS are causing a variety of problems for agricultural lands, roadsides, range lands, biodiversity, national parks, waterways, rivers, power dams, and urban green spaces, with serious economic and environmental significance (Gedyon Tamiru, 2017).

According to recent studies, new plant invaders that have not yet been discovered in Ethiopia are beginning to sprout, spread to uninvaded areas, and have harmful effects (Jemal Tola & Taye Tessema, 2015). While *E. plantagineum* was unheard of in the nation, emergent weeds like *Argemone mexicana, Senna didymobotrya*, and *Tagetes minuta* were discovered there.

## 2. Materials and Methods

### 2.1. Description of the Study Area

#### 2.1.1. Geographical location

The study was conducted in the Dejen woreda (district) of the Amhara regional state in Ethiopia. The study area was located in the East Gojjam zone of the Amhara regional state, on the edge of the Abay canyon. The geographical coordinates of the study area range from approximately 10°02′28′′N to 10°20′17′′N latitude and 38°02′30′′E to 38°18′27′′E longitude. The elevation of the area varies between 1002 and 2586 m.a.s.l. It is located at a distance of 230 km from Addis Ababa, 335km from Bahir Dar by Debre Markos, and 255km connected by Motta, which is the capital city of Amhara regional state. It is bordered on the South by Abay River which separates it from the Wara Jarso woreda in the North Shoa zone of the Oromia regional state, on the West by Awabel, on the Northwest by Debay Telatgen, on the North by Enemay, and on the East by Shebel Berenta woredas of East Gojjam zone of the Amhara regional state (Fig. 2).

**Figure 1:**
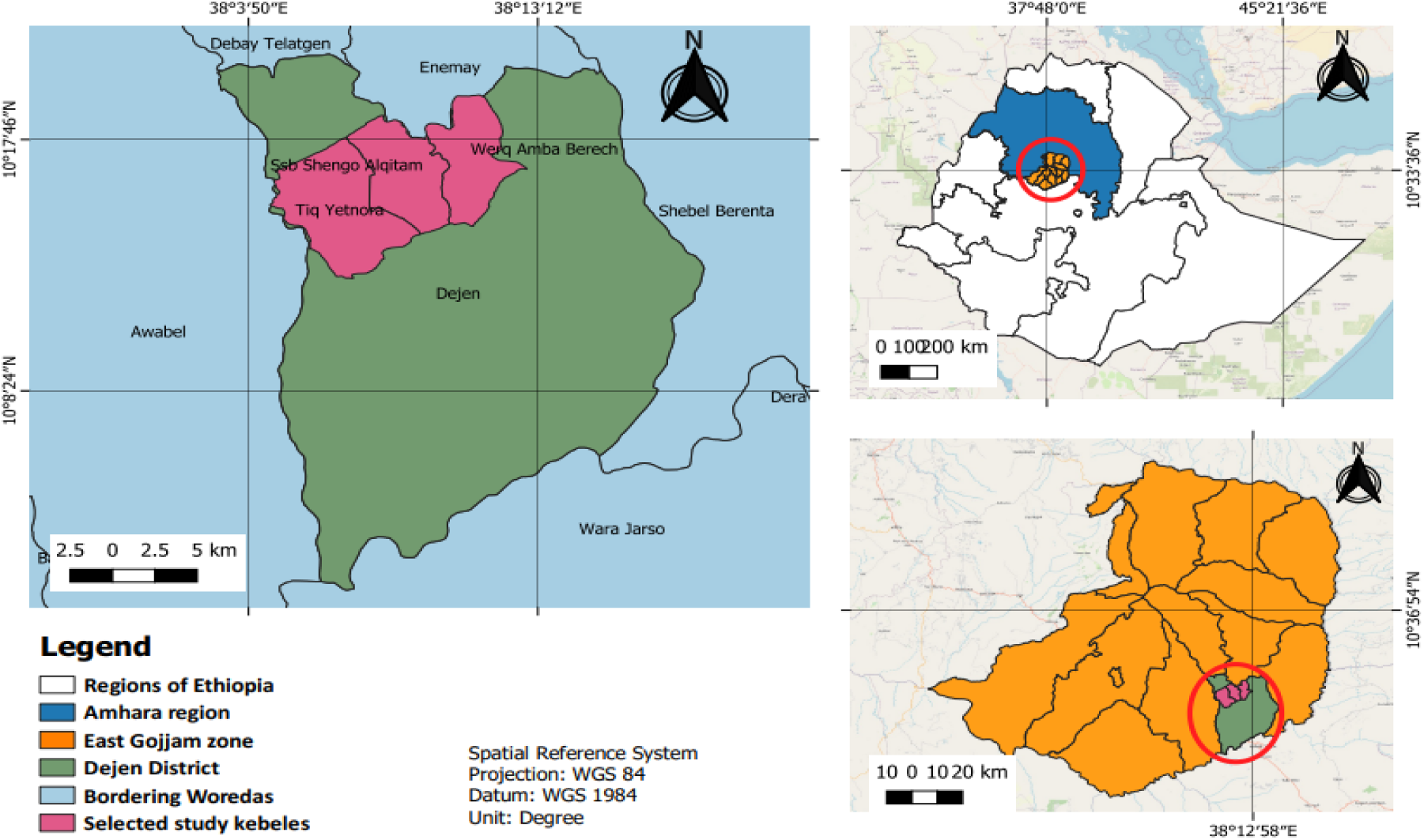
Location map of the study area.

**Figure 2:**
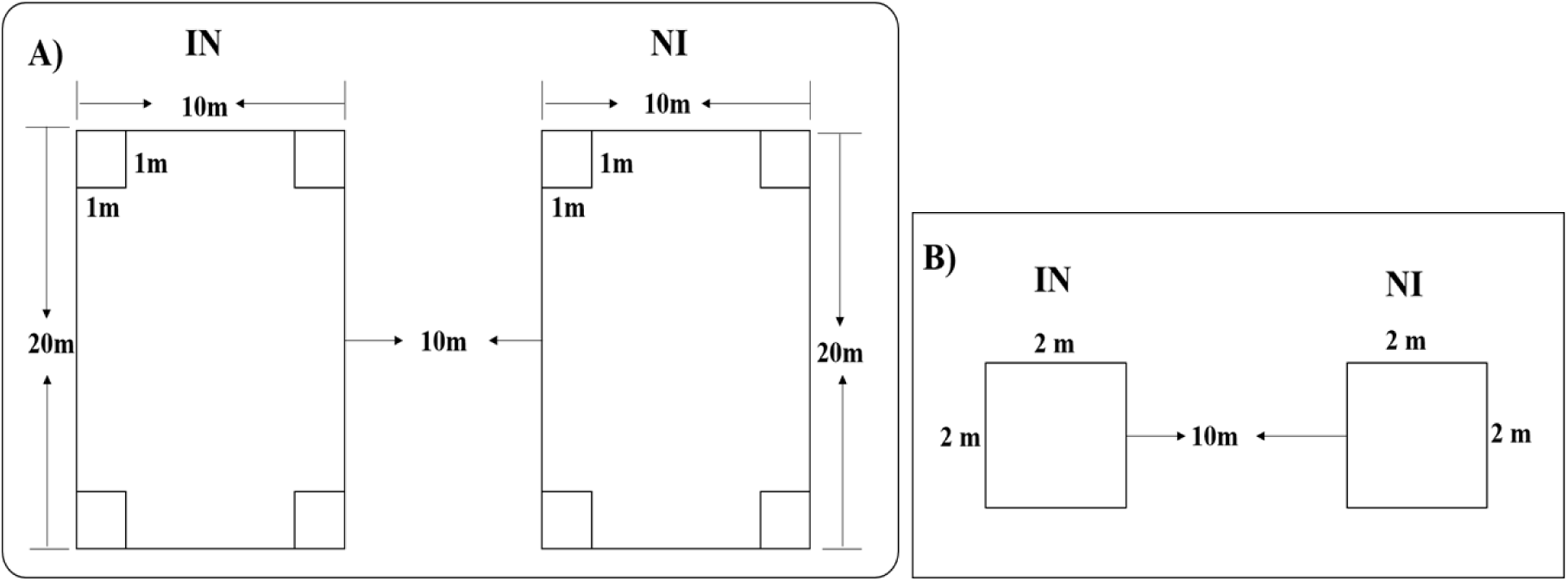
Layout of sample plots for plant and animal data collection (a) grazing land use type and (b) roadside land use type.

#### 2.1.2. Topography and climate

According to the information provided by the Dejen District Agriculture Office, the Dejen district has a diverse landscape consisting of plains, gorges, slopes, and hilly topography. Approximately 38.8% of the district is plains (highlands), 49.4% is categorized as Weyna Dega, and 11.8% is dipping (lowland). In terms of climate, Dejen experiences a moderate climate with temperatures ranging between 17°C (63°F) and 22°C (73°F) throughout the year. The mean annual maximum temperature is 27.44°C (81.39°F), while the mean annual minimum temperature is 15.37°C (59.67°F). The mean annual average temperature is around 23°C (73°F). The district receives an average annual precipitation of about 1341 mm (52.8 in.) spread across 148 rainy days annually, with a threshold of 1 mm (0.04 in.). Dejen enjoys an average of 3897 hours of sunshine per year, and the duration of daylight varies from 11 hours 31 minutes to 12 hours 41 minutes per day.

The district has a mean annual rainfall ranging from 800 mm to 1,200 mm. Approximately 90% of the district receives an average annual rainfall of 850 to 1,000 mm. Dejen has a unimodal rainfall pattern, with the rainy season typically extending from June to September. With an altitude range of 1,002 to 2,586 meters above sea level, the district’s climate and topography make it suitable for agricultural practices. The sustainable rainfall and moderate temperatures contribute to the agricultural productivity of the area (Dejen District Agricultural Office, 2022).

#### 2.1.3. Vegetation and soil type

According to the information provided by the Dejen District Agriculture Office (2022), the district has a variety of natural resources and soil types. The natural resources include natural forests, plantation trees, shrub land, bushland, and grassland, which contribute to the ecological diversity of the area. The major soil types in the district are vertisols (clayey soils with high clay mineral content), cambisols (weathered soils), nitisols (fertile soils used for agriculture), and entisols (minimal development of soil horizons) respectively in order of proportion from high to low in terms of area coverage.

Additionally, the district is known to have various types of rocks, such as limestone, sandstone, silica sand, calcite, marble, clay soil, and granite. These geological features indicate the presence of mineral resources in the area, which could potentially support industrial sectors. Therefore, due to its diverse soil types and geological composition, Dejen district has the potential to support both agricultural activities and industrial sectors, making it suitable for a range of economic activities (Dejen District Agricultural Office, 2022).

#### 2.1.4. Population

According to the 2022 national census conducted by the Central Statistical Agency of Ethiopia (CSA), Dejen woreda has a total population of 131,930. Out of this population, 63,715 are male and 68,215 are female. The woreda has a total area of 620.97 square kilometers or 62,097 hectares. With a population of 131,930, the population density of Dejen Woreda is approximately 121.5 people per square kilometer. In terms of religious affiliation, the majority of the inhabitants in Dejen woreda follow Ethiopian Orthodox Christianity, accounting for 97.01% of the population. Approximately 2.85% of the population identify as Muslims, while the remaining 0.14% follow other religions.

#### 2.1.5. Farming Systems and land uses

Dejen woreda is indeed known for its agricultural activities, with a focus on both crop cultivation and livestock rearing. The agricultural production system in the district is characterized as a mixed crop-livestock farming system. The majority of the population, over 98%, derives their livelihood from agricultural practices, including crop cultivation and animal-rearing activities, and the rest 2% or less do small business.

Cattle and goat production plays a central role in the lives of many households in Dejen woreda. Livestock rearing, alongside crop cultivation, contributes significantly to the local economy and sustains the livelihoods of the community.

In addition to its agricultural significance, Dejen woreda is home to various natural resources and landmarks. These natural resources, such as forests, rivers, and fertile lands, contribute to the agricultural productivity of the area. The district’s cultural and historical significance is also evident through its landmarks, which may include historical sites, traditional practices, and cultural events. Overall, the agricultural activities and natural resources in Dejen woreda play a crucial role in the lives of its residents, shaping their livelihoods and contributing to the local economy and cultural heritage of the region.

**Table 1:** Major land uses (Lu) and land cover (Lc) types in the Dejen district.

| № | Land Use and Land Cover Type | Area in hectare | % coverage |
| --- | --- | --- | --- |
| 1 | Forest | 3,522.51 | 5.7% |
| 2 | Shrubs | 2,422 | 3.9% |
| 3 | Cultivated lands | 37,023 | 59.6% |
| 4 | Grazing land | 1,980 | 3.2% |
| 5 | Non-use | 5,006 | 8% |
| 6 | Others | 12,143.49 | 19.6% |
| **Total** |  | **62,097** | **100%** |

Source: (Dejen District Agriculture Office, 2018)

### 2.2. Materials

The common equipment and tools used for plant specimen collection during the study included clippers (scissors), a digger, a field press (woody frame), straps, newspapers (blotters), pencils, cameras, a meter for laying out sample plots, and GPS tools for recording latitude, longitude, and altitude. Documentation data forms were also used, recorded in a notebook.

In the field, the materials used for plant data collection included GPS, a sampling frame, a digger, scissors, plastic bags, a camera, a notebook, a pencil, and tag labels. For herbarium presses, woody frames, straps, and blotters were utilized. For collecting invertebrates, a manually made insect net was used, along with small glass bottles for storing the collected insects. Needles were used to penetrate their bodies and attach them to boards made of different materials.

### 2.3. Methods

#### 2.3.1. Reconnaissance survey

A visit to the area was conducted to observe the distribution, spread, and effect of *E. plantagineum* on plant and animal species in different kebeles of the study area. The observation included assessing its presence in uncultivated areas, cultivated areas, along roadsides, grasslands (grazing land), and around construction sites. The purpose of the preliminary survey was to evaluate the distribution and level of infestation of this invasive species in various localities and land uses. The survey helped identify sites for detailed sampling of floral and faunal species and collection of these species. Based on this survey, sampling plots (quadrats) were established.

#### 2.3.2. Sampling design techniques

Based on the reconnaissance survey, the study sites were purposively selected based on the level and severity of infestation by *E. plantagineum*. For statistically sound sampling purpose infestation along roadsides and grazing lands were selected. These two dominant land use types (grazing land and roadside) invaded by *E. plantagineum* were selected using a stratified random sampling method from the three selected study kebeles. The reason for selecting these two areas is that *E. plantagineum* is highly distributed in these areas and has a significant impact. Based on these criteria, three kebeles were selected out of 22 kebeles, and from each level of infestation site (highly, moderately, and least) one kebele was selected (Table 3). For each kebele, five percent of the infested areas were sampled for both land use types (roadside and grazing land).

To collect data from grazing land use types, a 10m x 20m plot was established to study tree species, shrubs. Within this plot, four 1m x 1m subplots were located at each corner to collect data on herbaceous vegetation. On roadside land use types, a 2m x 2m plot was established to gather information on herbaceous vegetation, tree or shrub seedlings, (Fig. 3). This plot size was chosen since the size of the roadside land units is narrow. The distribution and infestation level of *E. plantagineum* invasions were assessed, and the number of sample plots was determined based on the area coverage of *E. plantagineum* invasives. More plots were sampled from highly infested areas, while fewer sample plots were taken from least infested areas (Appendix 2).

**Figure 3:**
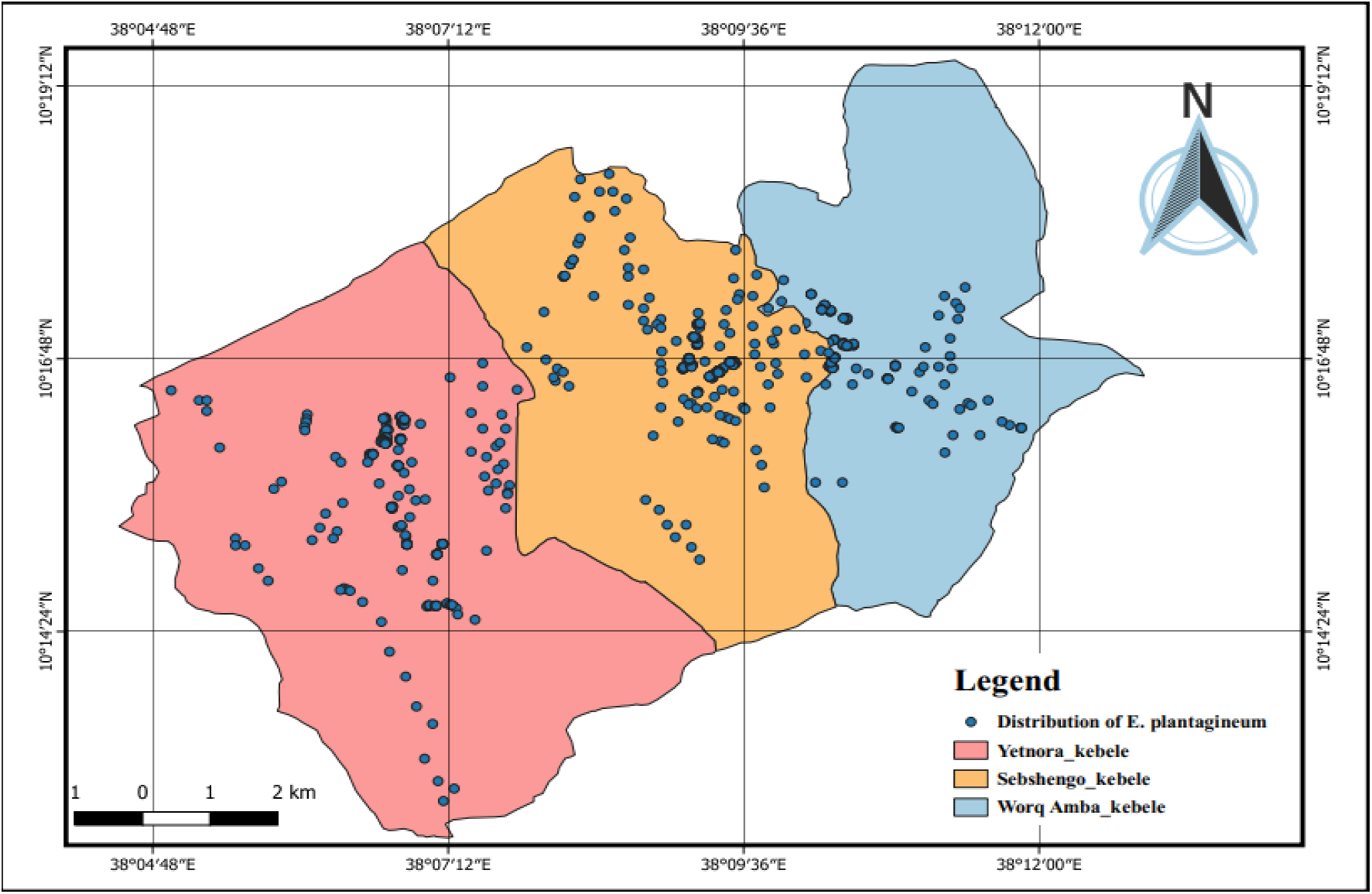
Distribution map of *Echium plantagineum* L. in three *Kebeles* of Dejen *Woreda*.

Thus, on both invaded and non-invaded roadside land use types twelve, ten, and six 2m x 2m plots (quadrats) were taken from Sebshengo, Yetnora, and Worq Amba kebeles respectively, whereas on grazing land use types (invaded and non-invaded plots) forty, thirty, and eighteen a 10m x 20m sample plots were taken from Sebshengo, Yetnora, and Worq Amba kebeles respectively (Table 3). Each of the 1m x 1m subplots and 2m x 2m plots were subdivided into a 10cm x 10cm grid to measure the percentage cover for each plant species (plant cover) (Junior *et al*., 2022). A Complete list of herbaceous plants was compiled for each quadrat, and the percent cover value was estimated for each species.

To examine the effects of *E. plantagineum* invasions on the floral species diversity of invaded communities, the layout of sampling plots was standardized across all three selected kebeles. The number of sample plots was equally distributed between invaded and adjacent non-invaded areas for both land use types. As much as possible data from non-invaded plots were collected within a distance of less than 10 m from invaded plots that exhibited similar ecological and environmental gradients (Karki, 2009).

The location (latitude, longitude, and altitude) of quadrats of the study site was recorded with the help of a GPS instrument (Amare Seifu & Ermias Lulekal, 2021). A total of 116 main plots were sampled, consisting of 58 invaded plots and 58 non-invaded plots, representing both land use types. Among these, 88 quadrats (44 plots for each invaded and non-invaded) were 10m x 20m in size, representing the grazing land use type. Within each of these main plots, 352 nested plots measuring 1m^2^ were placed at each corner. Additionally, 28 quadrats measuring 2m x 2m were sampled for the roadside land use type (Table 3 and Appendix 2).

**Table 2:**
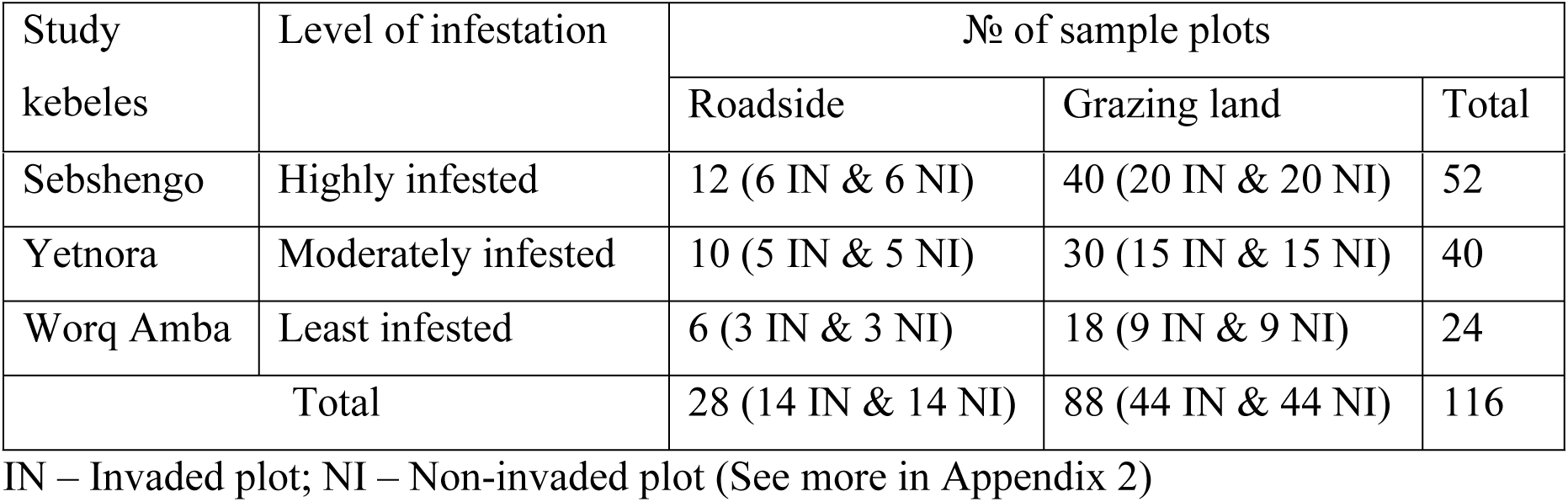
Numbers of quadrats for selected study sites (kebeles).

| Study kebeles | Level of infestation | No of sample plots |  |  |
| --- | --- | --- | --- | --- |
|  |  | Roadside | Grazing land | Total |
| Sebshengo | Highly infested | 12 (6 IN & 6 NI) | 40 (20 IN & 20 NI) | 52 |
| Yetnora | Moderately infested | 10 (5 IN & 5 NI) | 30 (15 IN & 15 NI) | 40 |
| Worq Amba | Least infested | 6 (3 IN & 3 NI) | 18 (9 IN & 9 NI) | 24 |
| Total |  | 28 (14 IN & 14 NI) | 88 (44 IN & 44 NI) | 116 |

IN – Invaded plot; NI – Non-invaded plot (See more in Appendix 2)

**Table 3:** Effect of *Echium plantagineum* L. on herbaceous and woody plant diversity, richness, and evenness.

| No | Parameters | Non-invaded | Invaded | % reduction over non-invaded |
| --- | --- | --- | --- | --- |
| 1 | Richness (total species) | 80 | 54 | (-) 32.5% |
| 2 | No. of Individuals | 832 | 552 | (-) 33.65% |
| 3 | Dominance_D | 0.047 | 0.578 | (+) 91.87% |
| 4 | Simpson_1-D | 0.953 | 0.422 | (-) 55.72% |
| 5 | Shannon_H' | 3.537 | 1.389 | (-) 60.73% |
| 6 | Evenness_e^H/S | 0.429 | 0.073 | (-) 83% |
| 7 | Jaccard's similarity index | 0.58 |  |  |
| 8 | Beta diversity | 0.42 |  |  |

(–) sign show less value and (+) sign show high value in invaded site

#### 2.3.3. Data collection

Data collection was conducted between January and February 2025 to assess the effect of *E. plantagineum* on floral species diversity in different infested land use types. Each native plant species, invasive plant species were identified and recorded. The percentage cover of the invasive and herbaceous species was visually estimated by subdividing plots into a 10cm x 10cm grid (Bibby, 1992).

Plant species discovered in the quadrats were identified in the field. Sample specimens of the recorded plant species were collected, pressed, properly dried, and taken to the Ethiopian Biodiversity Institute Herbarium for identification following (Amare Seifu & Ermias Lulekal, 2021). The collected specimens were identified by comparing them with authenticated specimens, consulting experts, and referring to the published volumes of Flora of Ethiopia and Eritrea (Hedberg, 1996; Mesfin Tadesse, 2004).

#### 2.3.4. Data analysis

Descriptive (frequency and percentage) and inferential statistical tools were used to present, interpret, and analyze the data. To determine the distribution of *E. plantagineum* invasive species, the study utilized QGIS 3.18.3 software. In this study, GPS coordinates or spatial polygons data indicating the invaded sites on the occurrence or presence of *E. plantagineum* at different locations within the study area was gathered. Using QGIS, this spatial data has imported into the software and georeferenced it to align with the study area’s underlying base map or reference system. This software was employed to generate a distribution map, providing visual representation and analysis of the invasive species’ spread and presence in the study area. The biophysical data, including the total number of plant species, and number of individuals were organized using Microsoft Office Excel. The data collected from the plot survey were analyzed using different statistical tools with the help of IBM SPSS (Statistical Package for Social Sciences) Statistics version 26 software.

In the study area, the diversity of species in *E. plantagineum*-infested sample sites was evaluated using several metrics, including the Shannon Diversity index (H’), species evenness (E), Dominance Index (D), and Simpson Index of Dominance (1-D). These metrics were analyzed using PAST software, which allowed for statistical analysis and interpretation of the diversity indices in relation to the presence and effect of *E. plantagineum*. The purpose was to evaluate the impact of *E. plantagineum* on the growth and distribution of other plant species.

The species evenness value ranges between 0 and 1. A value of 0 indicates that a few species are highly abundant, while a value of 1 signifies that all species are equally abundant in the sample. The index of dominance values range between 0 and 1. A value closer to 0 indicates a more diverse plot, while a value closer to 1 suggests that the plot is dominated by a few species. The pattern observed for dominance index values is opposite to that of Shannon index values. As the Shannon Diversity value increases, the Index of dominance value decreases.

To compare infested and non-infested areas Jaccard’s similarity index and beta diversity were calculated (Musese *et al*., 2020). The frequencies of each species in non-invaded and invaded sites were determined. Jaccard’s similarity index and Sorensen’s similarity index were computed to assess the similarity between non-invaded and invaded areas for the selected land use types in each sampling site (Karki, 2009).

Jaccard’s similarity index (JSI) was calculated as **JSI** = **c**/(**a** + **b** + **c**), where c represents the number of species common to both infested and non-infested areas, b denotes the species present in infested areas but absent in non-infested areas, and a represents the species present in non-infested areas but absent in infested areas. Alternatively, Jaccard’s similarity index can be calculated as **JSI** = **c**/(**a** + **b** ― **c**), where c is the number of species shared by invaded and non-invaded communities, a is the number of species in the invaded community, and b is the number of species in the non-invaded community. The coefficient ranges from 0 to 1, where a value of 1 indicates complete similarity and a value of 0 signifies complete dissimilarity between the compared areas.

According to Amare Seifu *et al*. (2017), the beta diversity index was calculated to assess the impact of invasion on species composition and the relative covers of resident species in invaded and non-invaded study sites. The calculation of the Beta diversity index was based on species covers (Hejda *et al*., 2009). It is calculated using the formula **β*=*1 - JSI,** where JSI is Jaccard’s similarity index (Baselga, 2010).

A linear regression and correlation analysis was conducted to evaluate the relationship between the abundance of *E. plantagineum* and the abundance of floral and species within study quadrats. The regression equation was computed as **y** = **a** + **bx**, where y represents the dependent variable (parameters), a is the regression constant, b is the estimated regression coefficient, and x represents the independent variable (parameter). In this case, the independent variable (x) is the abundance of *E. plantagineum*, while the dependent variable (y) is the abundance of specific dominant non-invasive species.

## 3. Results and Discussion

### 3.1. Distribution of *Echium plantagineum* L. in the Study Area

*Echium plantagineum* L. has been observed to grow on roadsides, grazing land, and occasionally on poorly tilled farmlands. Among these habitats, roadside and grazing land use types are frequently infested by this invasive weed (Appendix 1). The high frequency of infestation along roadsides can be attributed to continuous disturbance and the transportation of sand and soil during road construction and maintenance. The distribution of *E. plantagineum* is expanding rapidly in the research area due to its prolific seed production, posing a significant threat to roadsides and grazing lands.

A similar study was conducted on the abundance and geographical distribution of invasive weed species in the Western Amhara Region, Ethiopia, by Melkamu Birhanie *et al*. (2020), who reported that the distribution of *E. plantagineum* is escalating at an alarming rate and invading roadsides and grazing lands along the Dejen to Debre Markos Road. Among the three selected study kebeles, Sebshengo kebele exhibited the widest distribution of *E. plantagineum* compared to the other two kebeles (Fig.4). The invasive species has covered substantial portion of the areas within the kebeles. In contrast, Worq Amba kebele showed the least colonization by *E. plantagineum* invasives (Fig. 4).

**Figure 4:**
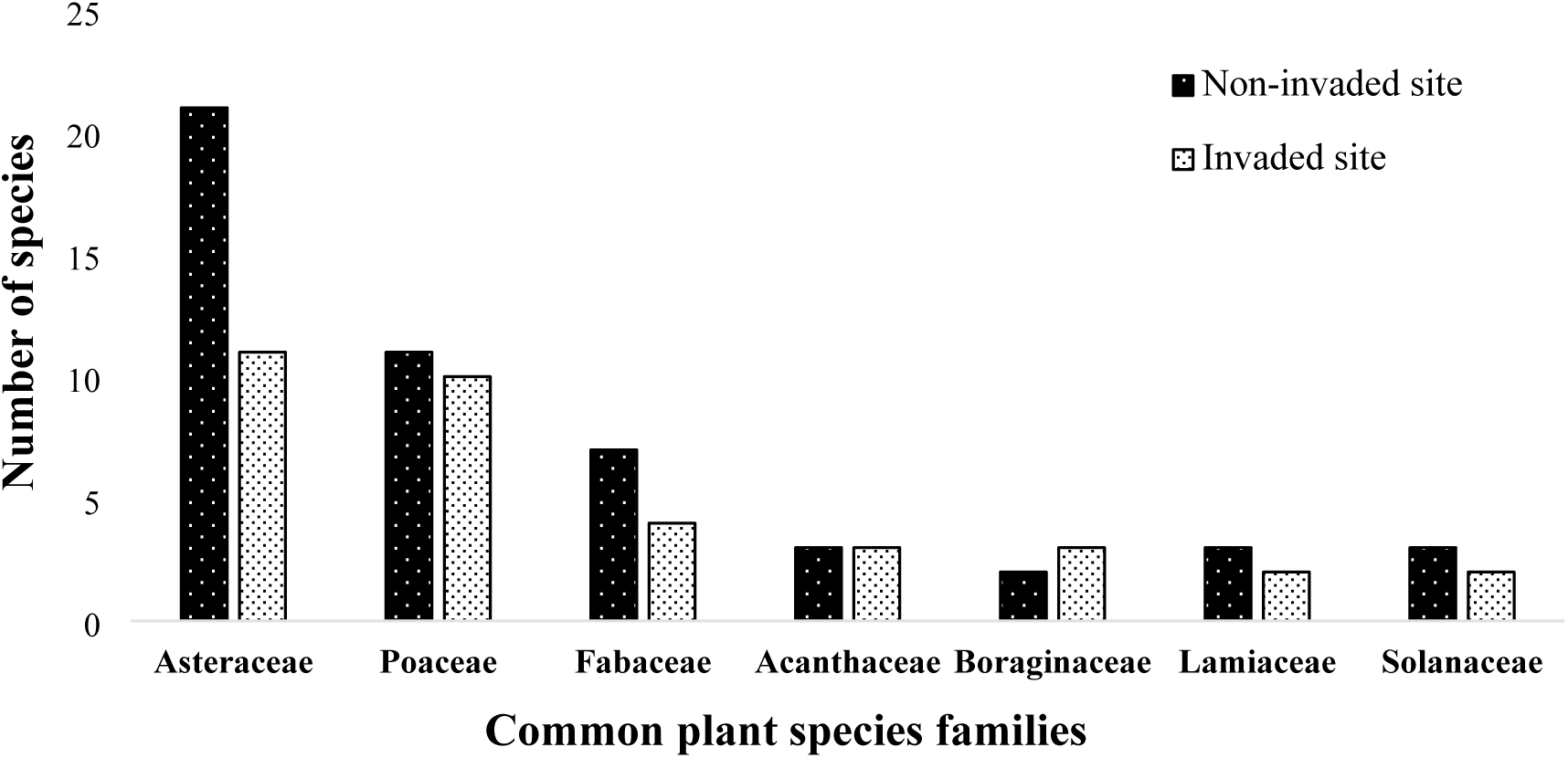
Comparison of plant families between invaded and non-invaded plots.

### 3.2. Effect of *Echium plantagineum* L. on Plant Species Diversity

In the non-invaded area, there were a record of 80 plant species belonging to 30 families. Whereas, in the *E. plantagineum* invaded sites, there were a record of 54 plant species belonging to 24 families (Appendix 3). Among 85 plant species, 49 species were found common in non-invaded and invaded areas. Thirty-one plant species (*Acacia mearnsii* De Wild., *Achyranthes aspera* L., *Acmella caulirhiza* Del., *Argemone mexicana* L., *Artemisia ludoviciana* Nutt., *Calpurina aurea* (Ait.) Benth., *Cirsium vulgare* (Savi) Ten., *Clematis hirsuta* Perr. & Guill., *Crepis dioscoridis* L., *Cucumis ficifolius* A.Rich, *Desmodium intortum* (Mill.) Urb., *Dodonaea viscosa* (L.) Jacq., *Erigeron bonariensis* L., *Ficus palmata* Forssk., *Laggera crispata* (Vahl) Hepper & J.R.I.Wood., *Lippia adoensis* Hochst. ex Walp., *Leucas martinicensis* (Jacq.) R.Br., *Nuxia congesta* Fresen., *Osyris lanceolata* Hochst. & Steud., *Premna schimperi* Engl., *Rhus vulgaris* Meikle, *Rhynchosia ferruginea* A. Rich., *Rosa abyssinica* Lindley, *Sonchus asper* (L.) Hill, *Solanum stramoniifolium* Jacq., *Sphaeranthus suaveolens* (Forssk) DC., *Stephania abyssinica* (Dill. & Rich.) Walp., *Tagetes minuta* L., *Verbascum benthamianum* L., *Verbena officinalis* L., and *Vernonia amygdalina* Del.) were found only in non-invaded areas. While five species (*Cupressus lusitanica* Mill., *Echium plantagineum* L., *Euphorbia candelabrum* Trema., *Sesbania sesban* (L.) Merr., and *Maytenus arbutifolia*. (A. Rich) Wilezek.) were growing only in the invaded areas.

Thus, the number of species and the number of families were reduced by 32.5% and 20% respectively in the *E. plantagineum-*invaded area as compared to the non-invaded area. In addition, the number of individuals also reduced by 33.65% in the invaded area compared to non-invaded. The decline in plant species and families indicates that the *E. plantagineum* invasion could have resulted in a reduction in plant diversity and a change in the composition of plant communities in the invaded sites. The results of the study indicate that the invasion of *E. plantagineum* had an impact on the diversity and composition of flora in the study area.

The Shannon Diversity index value for non-invaded plots was 3.537, while for invaded plots it was 1.389. Similarly, the species evenness index value for non-invaded plots was 0.429, while for invaded plots it was 0.073. Thus, Shannon’s index of diversity and evenness index value were reduced by 60.73% and 83% respectively in the *E. plantagineum* invaded areas compared to non-invaded plots (Table 4). These findings indicate that the invasion of *E. plantagineum* has resulted in a decrease in herbaceous and woody plant species richness, diversity, and evenness. These results showed that the non-invaded areas have a higher Shannon Diversity index (3.537) and higher species evenness value (0.429) compared to the *E. plantagineum* invaded areas. The lower values of Shannon’s index of diversity (1.389) and lower evenness index value (0.073) suggest that the invaded communities have a lower variety of species. The distribution of individuals among species is uneven, with one or a few species dominating the invaded areas. Evenness has values between 0 and 1, where 1 represents a situation in which all species are equally abundant.

**Table 4:** Pearson correlations between the abundance of *Echium plantagineum* L. and plant species abundance.

| Correlations |  | *C. dactylon* | *A. distachyos* | *C. longus* |
| --- | --- | --- | --- | --- |
| *E. plantagineum* | Pearson Correlation | -.291** | -.319** | -.251** |
|  | Sig. (2-tailed) | .002 | .000 | .007 |
|  | N | 116 | 116 | 116 |

\*\* . Correlation is significant at the 0.01 level (2-tailed), and N – No of plots.

This finding is similar to the findings of studies by Dogra *et al*. (2009), Amare Seifu *et al*. (2017), Byabasaija *et al*. (2020), and Amare Seifu & Ermias Lulekal (2021), which found that the lower values of Shannon’s index of diversity (H’), and evenness index value in the invaded areas. In contrast, the higher values of Shannon’s index of diversity and evenness index in non-invaded communities indicate greater species heterogeneity and a more balanced distribution of individuals among different species. Accordingly, a lower value of the Shannon Diversity index suggests an area was dominated by a few species, i.e., invaded sites were relatively homogeneous species in the community. Therefore, in the non-invaded study sites, the communities were more diverse in the type of species and heterogeneous in the community.

According to Barnes (1998), invaded communities have low diversity because invasive plants alter the invaded ecosystem and species composition to such an extent that they threaten native flora. Thus, the higher value of the Shannon Diversity index in the non-invaded plots suggests that *E. plantagineum* has an impact on the indigenous species composition through a decrease in species diversity. This harmonizes with many studies conducted in different parts and vegetation types of the world (Hailu Shiferaw *et al*., 2003; Melkamu Kifetew & Zerihun Woldu, 2021), which have reported that in natural areas the shrub has a series of deleterious effect on some plant species and is known to displace natural bush communities as well as prevent natural regeneration of some tree species.

The Jaccard’s similarity index of the communities in the invaded and non-invaded areas was 58% (similarity index), which showed loss of species was due to the invasion of *E. plantagineum* and that of the beta diversity index (dissimilarity index) was 42%. This indicated that *E. plantagineum* invaded site is dominated by a limited number of species. The uneven distribution of individuals among species in the invaded areas suggests that only a small number of species are well-adapted to the environment altered by *E. plantagineum* invasion. If those few species that can survive are extremely affected, then the site that contains the species may be disturbed as compared to non-invaded areas.

This finding is consistent with the Amare Seifu *et al*. (2017) study, which reported that a low diversity index value suggests an area is dominated by a limited number of plant species. The dissimilarity values showed significant differences between *E. plantagineum* invaded and non-invaded plots. Similarly, this finding is in agreement with Kohli *et al*. (2004) and Mesfin Boja *et al*. (2022), who reported fact that plant species diversity in the non-invaded area was greater than in the *E. plantagineum*-invaded areas. Furthermore, the index of dominance (D) value was found to have increased by 91.87% in the invaded areas compared to the non-invaded areas (Table 4). The increase in dominance indicates that a smaller number of plant species have become more dominant in the invaded areas, potentially leading to a shift in the composition and structure of the plant community.

This finding is in line with the studies by Dogra *et al*. (2009), Wambua (2010), Gebrehiwot & Berhanu (2015), Amare Seifu & Ermias Lulekal (2021), and Melkamu Kifetew & Zerihun Woldu (2021) who found that a higher index of dominance value in invaded areas and a lower index of dominance value for floral communities in non-invaded areas would indicate that the invaded communities were homogenous whereas non-invaded floral communities would be indicated that the heterogeneous species nature. These results collectively highlight the negative impact of *E. plantagineum* invasion on plant diversity and community dynamics in both herbaceous and woody vegetation.

#### 3.2.1. Effect of *Echium plantagineum* L. on plant species composition

The results of this study showed that the invasion of *E. plantagineum* affected the composition of woody and herbaceous plant species in the study area. At the family level, in the non-invaded areas, of the 30 families, the family Asteraceae represented the highest number of species (21 species), accounting for 26.3%, followed by Poaceae (13.8%), and Fabaceae (8.8%). It is worth noting that the above-mentioned families alone represent 48.9% of plant species in the total flora in non-invaded study sites (Appendix 3). Whereas, in the invaded areas, of the 24 plant families, the Asteraceae family was the highest-ranking in the number of species (11 species) accounting for 20.4%, followed by Poaceae (18.5%), and Fabaceae (7.4%). The above-mentioned three families shared 46.3% of plant species recorded in the *E. plantagineum*-invaded study areas (Appendix 3).

The current study indicated that the three families Asteraceae, Poaceae, and Fabaceae were the dominant families for the whole identified plant species for the study sites (Fig. 5). This finding is in agreement with Tsegay Gebregergs (2019), who studied herbaceous species composition which indicated that the three families Poaceae, Asteraceae, and Fabaceae took the lead of the herbaceous species sampled. Further studies by Lema Etefa (2011), Genet Atsbeha *et al*. (2015), Amare Seifu *et al*. (2017), and Bhatta *et al*. (2020) on the floristic composition of herbaceous flowering plant species found the three families Poaceae, Asteraceae, and Fabaceae have the highest number of species.

**Figure 5:**
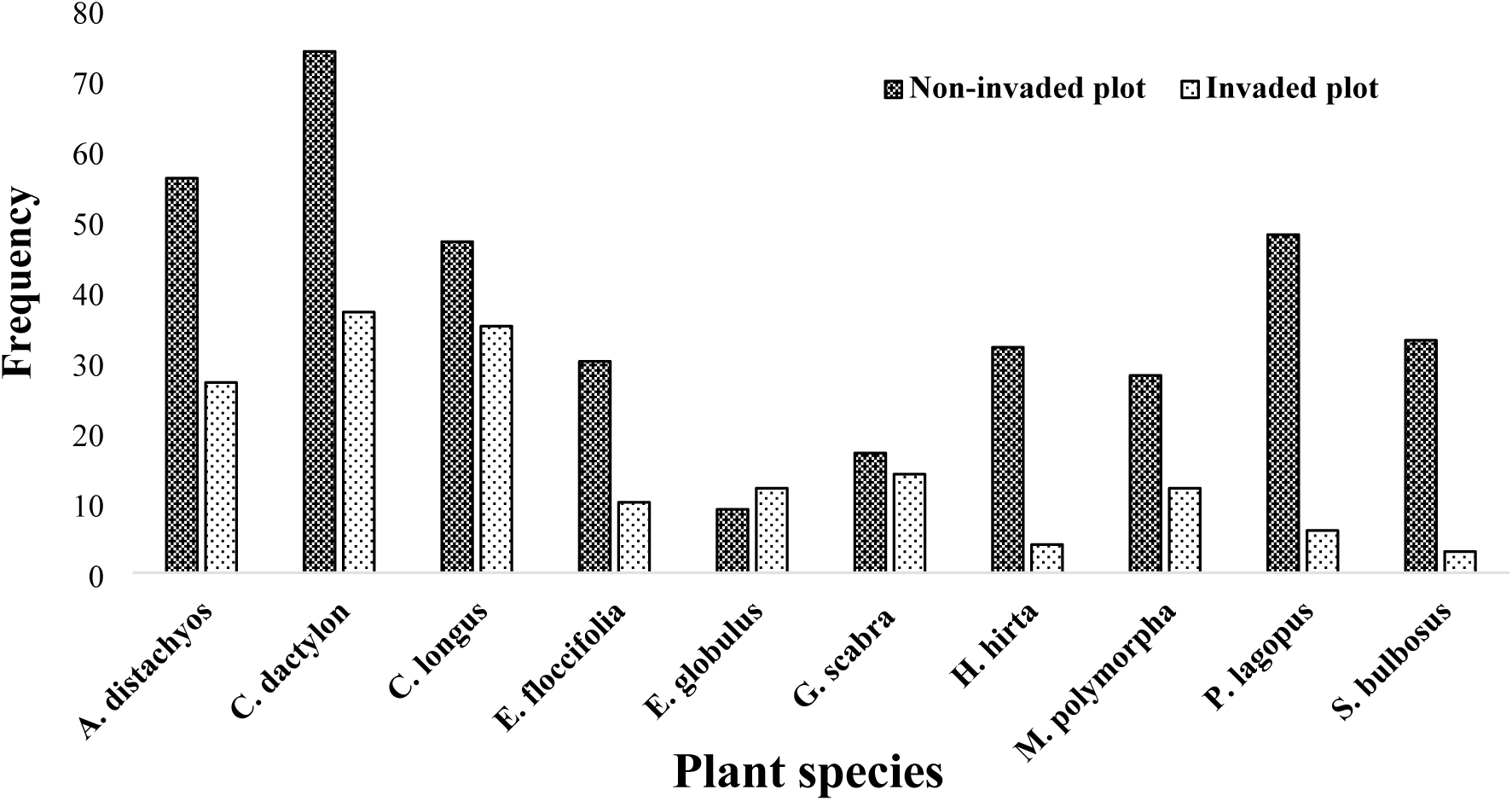
Comparison of some frequently recorded plant species between invaded and non-invaded plots.

Out of the 80 plant species recorded from non-invaded plots, *Cynodon dactylon* had the highest frequency, followed by *Andropogon distachyos*, *Plantago lagopus*, and *Cyperus longus* (Fig. 6). On the other hand, out of the 54 plant species recorded from invaded plots, *Echium plantagineum* had the highest frequency, followed by *Cynodon dactylon*, *Cyperus longus*, and *Andropogon distachyos* (Fig. 6).

**Figure 6:**
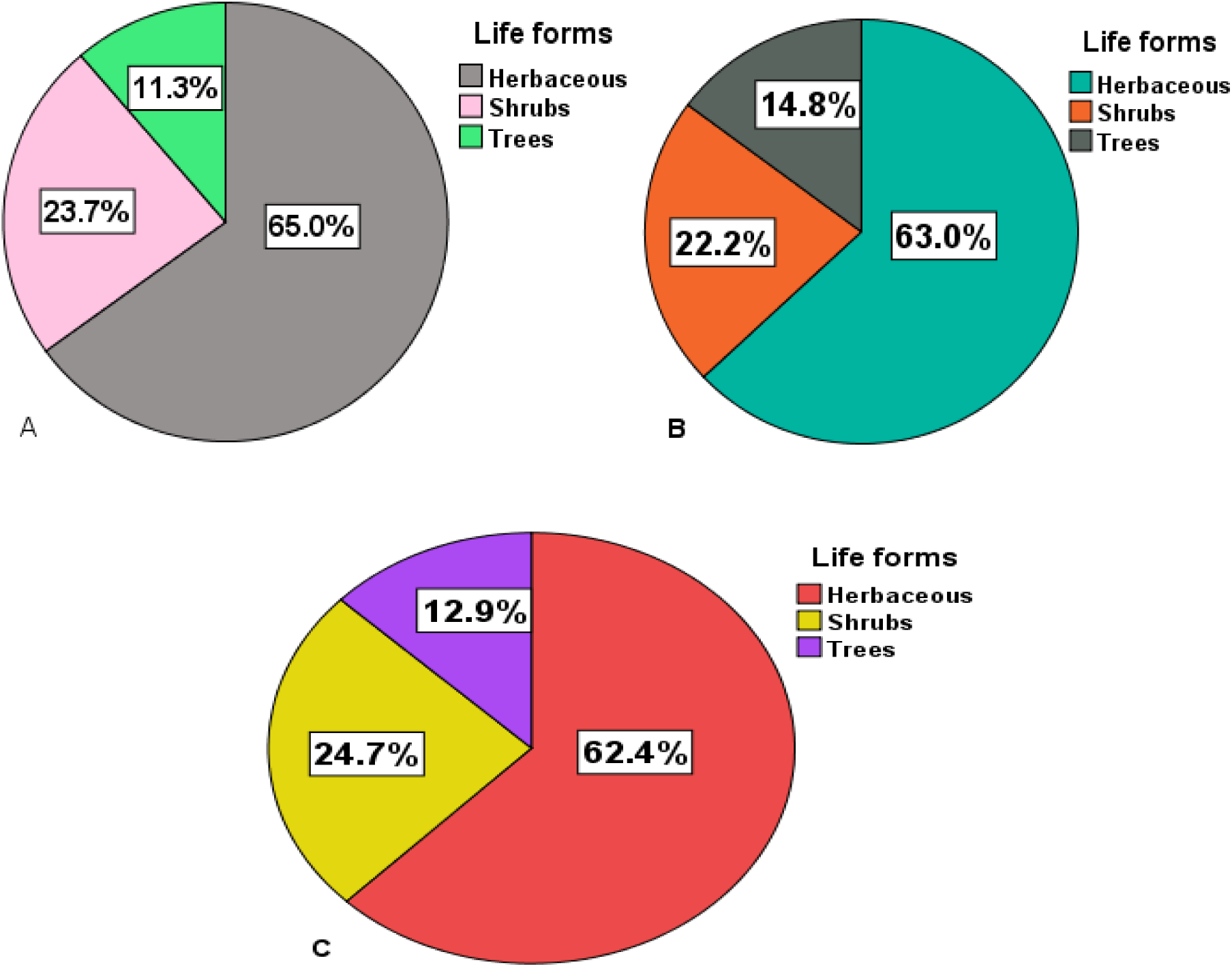
Different life forms of plant species recorded (a) non-invaded plot, (b) invaded plot, and (c) entire plot.

Regarding to growth habits, among 80 plant species identified from non-invaded plots, the majority were herbaceous species (52 species), followed by shrub species (19), and nine tree species (Fig. 7a). Whereas, among the 54 plant species identified from invaded plots, 34 species were herbaceous, followed by shrub species (12), and eight tree species (Fig. 7b). Overall, a total of 85 documented plant species, including 53 herbaceous species, 21 shrub species, and 11 tree species, were recorded from the entire study sites of both land use types (Fig. 7c).

**Figure 7:**
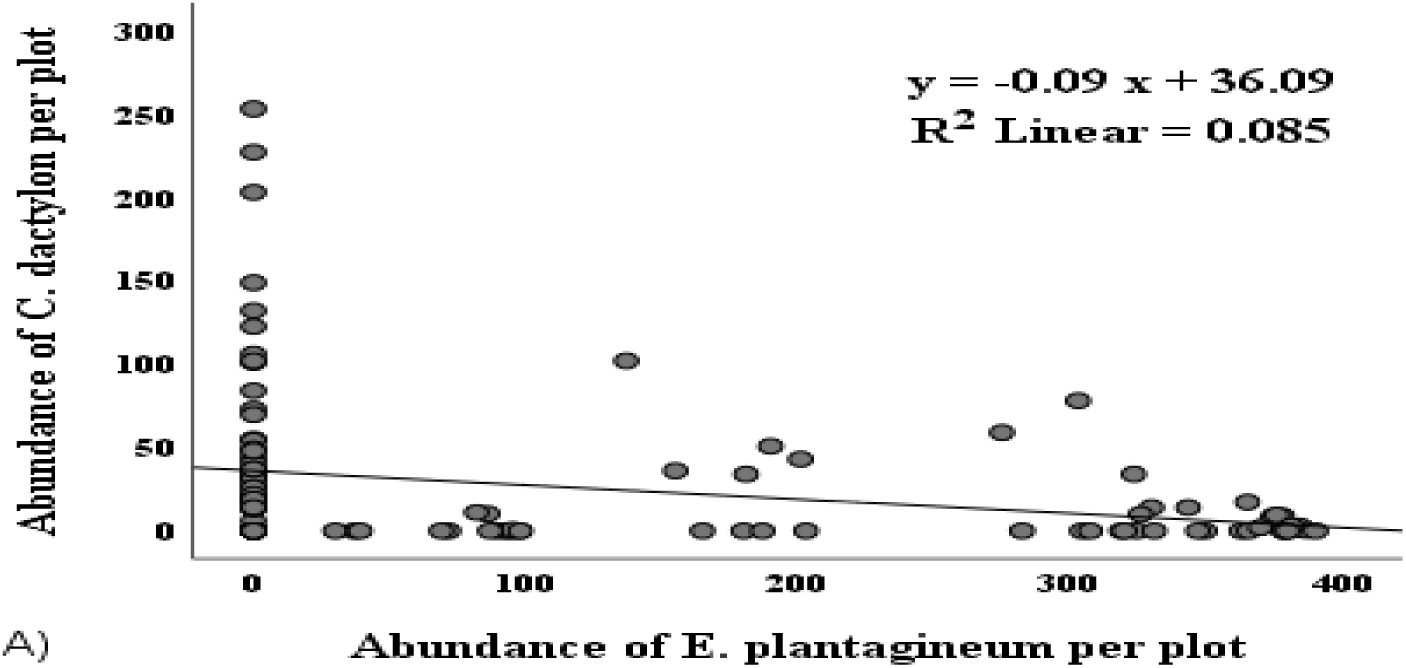
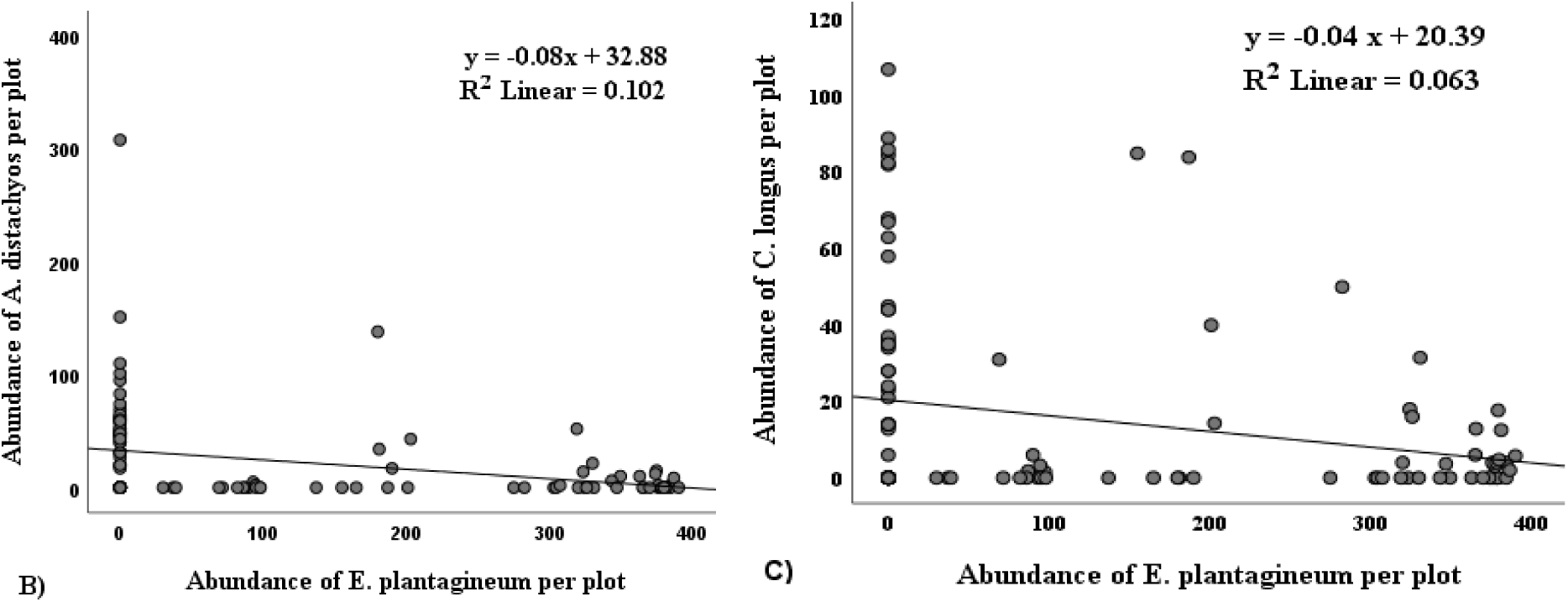
Relationship between the abundance of *E. plantagineum* and abundance of plant species (a) abundance of *E. plantagineum* Vs abundance of *C. dactylon* (b) abundance of *E. plantagineum* Vs abundance of *A. distachyos* and (c) abundance of *E. plantagineum* Vs abundance of *C. longus* per plot.

A pie chart (Fig. 7) depicted that the majority of plant species recorded from both invaded and non-invaded plots were herbaceous species, surpassing the number of shrub or tree species.

#### 3.2.2. Relationship between the abundance of *Echium plantagineum* L. and the abundance of non-invasive plant species

In all of the research plots, *Cynodon dactylon, Andropogon distachyos*, and *Cyperus longus* were the three most abundant native plant species. A scatter plot revealed that an increase in the abundance of *E. plantagineum* led to a decrease in plant species’ abundance of native flora. The results indicated that there was a negative linear relationship between the abundance of *E. plantagineum* and the abundance of *C. dactylon* species in each study plot. Thus, the regression equation can be presented as **y = -0.09 x + 36.06**, where y is an abundance of *C. dactylon* and x is the abundance of *E. plantagineum* per plot. In this case, R^2^ = 0.085, or 8.5% indicating that there is a weak negative relationship between the abundance of *E. plantagineum* and the abundance of *C. dactylon* plant species (Fig. 8a). The regression equation and Pearson correlation (r = -0.291, p = 0.002) at p = 0.01 indicated the existence of a negative linear relationship between the abundance of *E. plantagineum* and the abundance of *C. dactylon* species in each plot.

The results showed that there was also a negative linear relationship between the abundance of *E. plantagineum* and the abundance of *A. distachyos* plant species in the study plot. Thus, the regression equation can be written as **y = -0.08 x + 32.88**, where y represents the abundance of *A. distachyos* and x is the abundance of *E. plantagineum* per plot. In this case, R^2^ = 0.102, or 10.2% indicated that there is a weak negative relationship between the abundance of *E. plantagineum* and the abundance of *A. distachyos* plant species (Fig. 8b). The regression equation and Pearson correlation (r = -0.319, p = 0.00) at p = 0.01 indicated the existence of a negative linear relationship between the abundance of *E. plantagineum* and the abundance of *A. distachyos* species per plot (Table 6).

The regression analysis also showed a negative relationship between the abundance of *E. plantagineum* and *C. longus* species abundance per plot. Hence, the regression equation can be presented as **y = -0.04 x + 20.39,** where y is *C. longus* abundance and x is *E. plantagineum* abundance per plot. The regression equation indicated that there was a weak negative relationship between the abundance of *E. plantagineum* and *C. longus* species abundance. As the abundance of *E. plantagineum* increased species abundance of *C. longus* decreased (Fig. 8c). In this case, R^2^ = 0.063 or 6.3% of the dependent variable (*C. longus* species abundance) can be explained by the independent variable (abundance of *E. plantagineum*). The regression equation and Pearson correlation (r = -0.251, p = 0.007) at p = 0.01 indicated the existence of a negative linear relationship between the abundance of *E. plantagineum* and the abundance of *C. longus* species per plot.

All scatter plots (Fig. 8) showed that there was a negative linear relationship between the abundance of *E. plantagineum* and the abundance of native plants. The regression analyses revealed that as the abundance of *E. plantagineum* increased, the abundance of *C. dactylon*, *A. distachyos*, and *C. longus* decreased. The R^2^ values indicated that only a small percentage of the variation in native plant abundance could be explained by the abundance of *E. plantagineum*. Overall, these findings suggest that *E. plantagineum* has a negative influence on the abundance of native plant species in the studied plots.

Multiple studies consistently demonstrate that invasive plants have negative effects on native plant communities. These effects include reductions in abundance, diversity, richness, and evenness of native plant species. The findings from Rejmanek & Richardson (1996), D’Antonio & Meyerson (2002), Daehler (2003), Vilà *et al*. (2011), Catford *et al*. (2012), and Pyšek *et al*. (2012) collectively support the notion that invasive plants outcompete native species for resources, leading to declines in native plant abundance and diversity. Specifically, observed the negative relationship between the abundance of the invasive plant species *E. plantagineum* and the abundance of native plant species *C. longus*, *C. dactylon*, and *A. distachyos* suggests that the presence of *E. plantagineum* may indeed negatively affect the abundance of these native plants.

## 4. Conclusion and Recommendations

### 4.1. Conclusion

*Echium plantagineum* L. was primarily observed in Yetnora Kebele, which serves as the location for the Kajima Road construction contractor camp. It is concerning that its distribution is rapidly increasing, invading roadsides and grazing lands within the study area. The high frequency of its presence along roadsides can be attributed to road construction and maintenance activities. The result of the study indicates that *E. plantagineum* is the dominant invasive alien plant species (IAPS) in the selected study Kebeles within the study area. Its presence has a significant impact on the diversity and composition of both plant and animal species in the invaded communities. The vegetation in invaded areas showed lower species richness and evenness of herbaceous and woody plant species compared to non-infested vegetation in both land use types. Further, the infestation of invasive also had negative effects on animal species diversity and composition. This finding reported that higher diversity indices were relatively recorded in the non-invaded areas in all study sites when compared to invaded areas. The study findings indicated a negative relationship between the abundance of *E. plantagineum* and the abundance of faunal and floral species diversity. This suggests that as the abundance of *E. plantagineum* increased, the diversity of faunal and floral species decreased. Overall, these findings highlight the negative effects of *E. plantagineum* on the diversity and composition of both plant and animal species in the invaded areas.

### 4.2. Recommendations

Based on the results obtained from the study, several recommendations were put forward:

✓ Proper management strategies and control measures are necessary to prevent further spread and mitigate its negative ecological and economic consequences.
✓ Create public awareness about the negative impacts of *E. plantagineum* on plant and animal productivity, and biodiversity, especially among farmers. This will aid in timely and effective management, preventing further spread into new areas.
✓ Identify priority kebeles in the district to control the spread of *E. plantagineum*. Targeted measures can be implemented in these areas to prevent its expansion.
✓ Protect and restore sensitive land use types through an integrated approach, considering ecological needs. Various strategies should be employed to manage *E. plantagineum* in these areas.
✓ Conduct further studies to assess the impacts on crop yield, human and animal health, and biodiversity. These studies will inform management practices in the study area.
✓ Establish communication links between agricultural offices and farmers at different levels to detect and eradicate the spread and negative impacts of *E. plantagineum*. This coordination will enable effective response and management.

## Appendices

## Appendix 1: Sample plot (quadrat) calculation

| Selected kebeles | Level of infestation | Area description in hectare |  |  |  |  |  |  |  | № of plots (IN) |  |  |
| --- | --- | --- | --- | --- | --- | --- | --- | --- | --- | --- | --- | --- |
|  |  | Total | CL | GL | Others | IN GL | Sampled (5%) | IN RS | Sampled (5%) | GL plot | RS plot | Total |
| Sebshengo | High | 2956.5 | 2543.2 | 162.3 | 251 | 8.09 | 0.4045 | 0.051 | 0.0026 | 20 | 6 | 26 |
| Yetnora | Moderate | 4400.8 | 3520.8 | 187.48 | 692.52 | 5.95 | 0.2975 | 0.042 | 0.0021 | 15 | 5 | 20 |
| Worq Amba | Least | 2777.7 | 2252 | 135.25 | 390.45 | 3.48 | 0.174 | 0.025 | 0.0013 | 9 | 3 | 12 |
| Total no. of plots taken from both invaded LUT |  |  |  |  |  |  |  |  |  |  |  | 58 + 58 |

## Appendix 2: Total number of identified plant species in the study

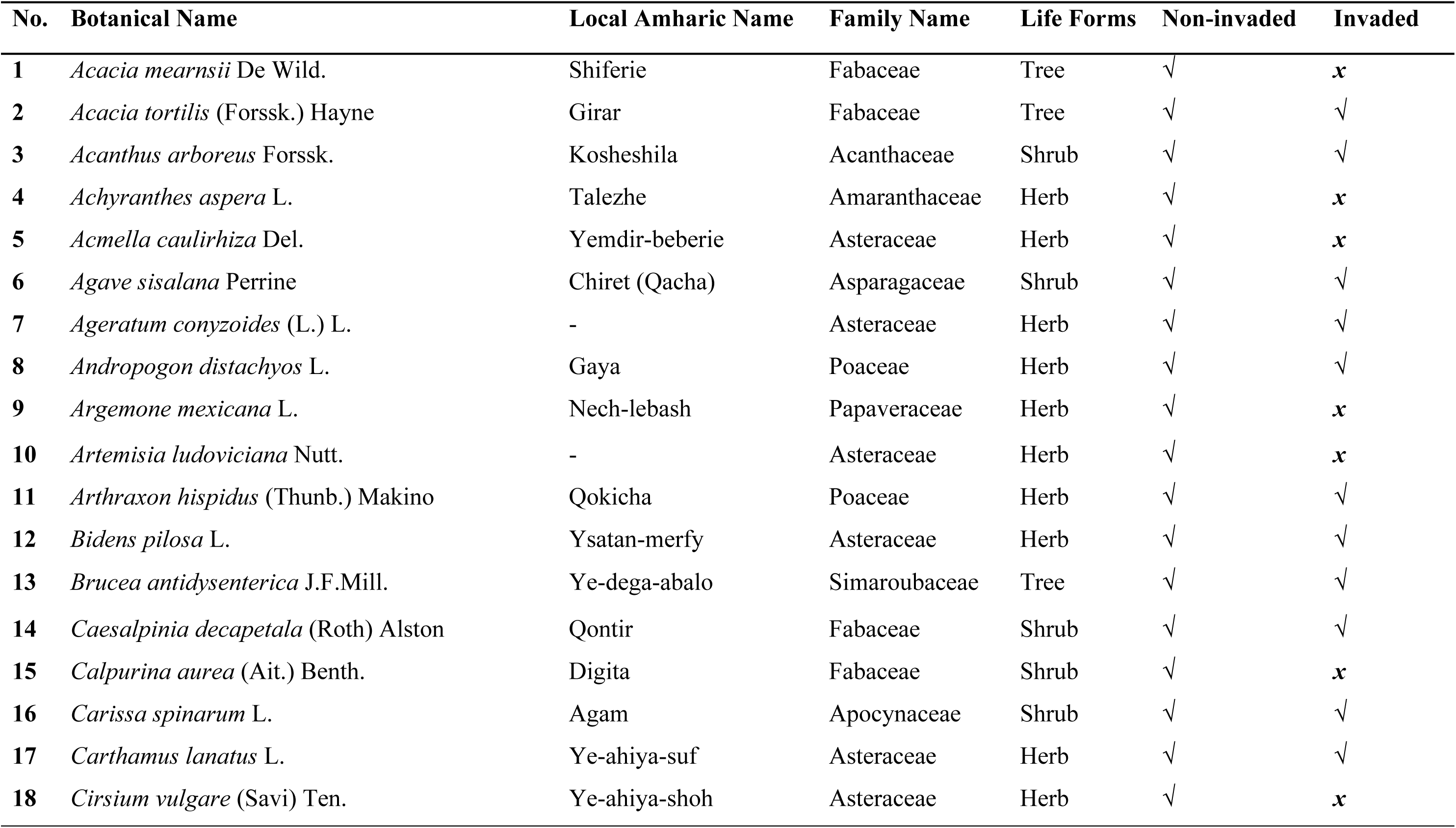
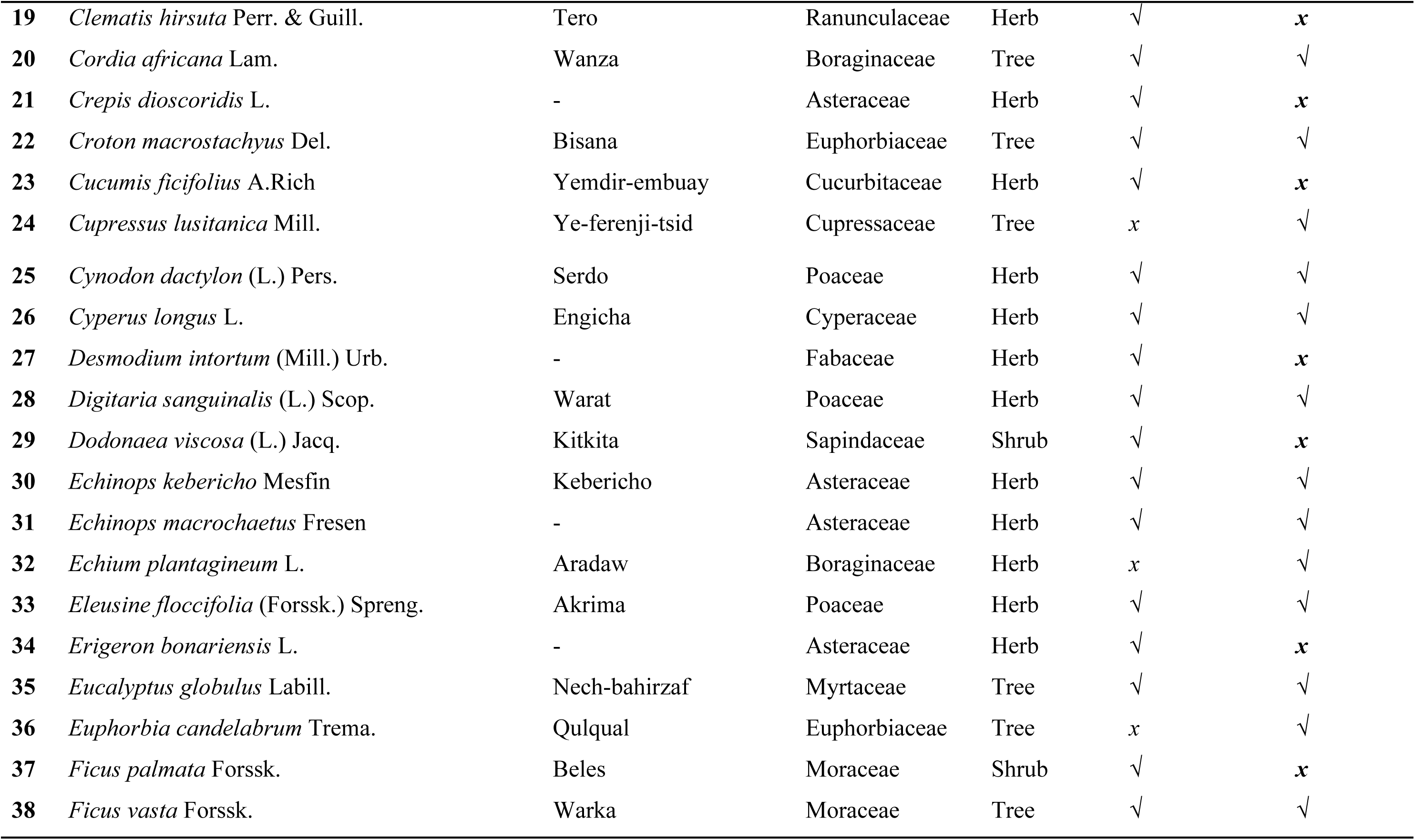
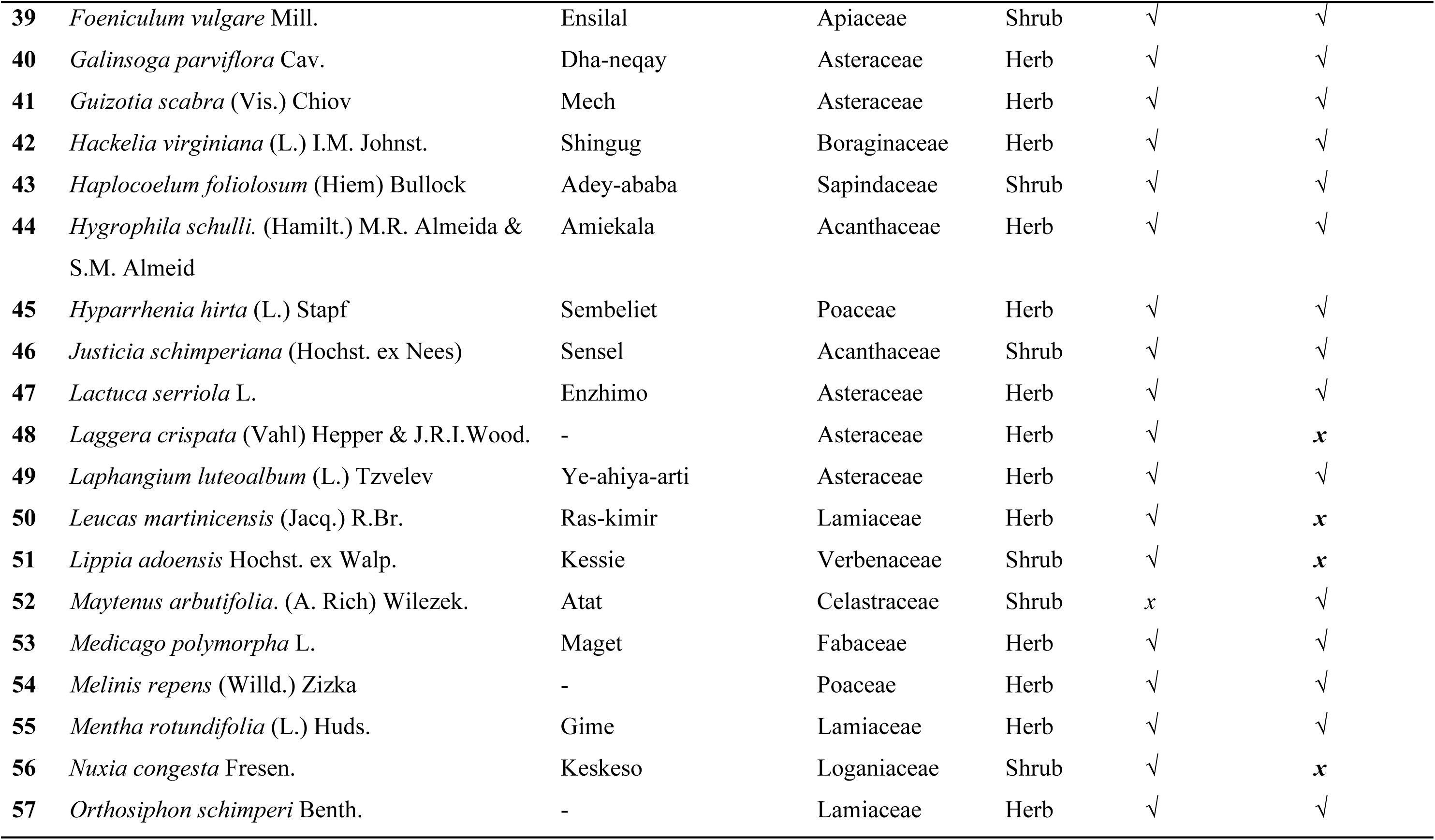
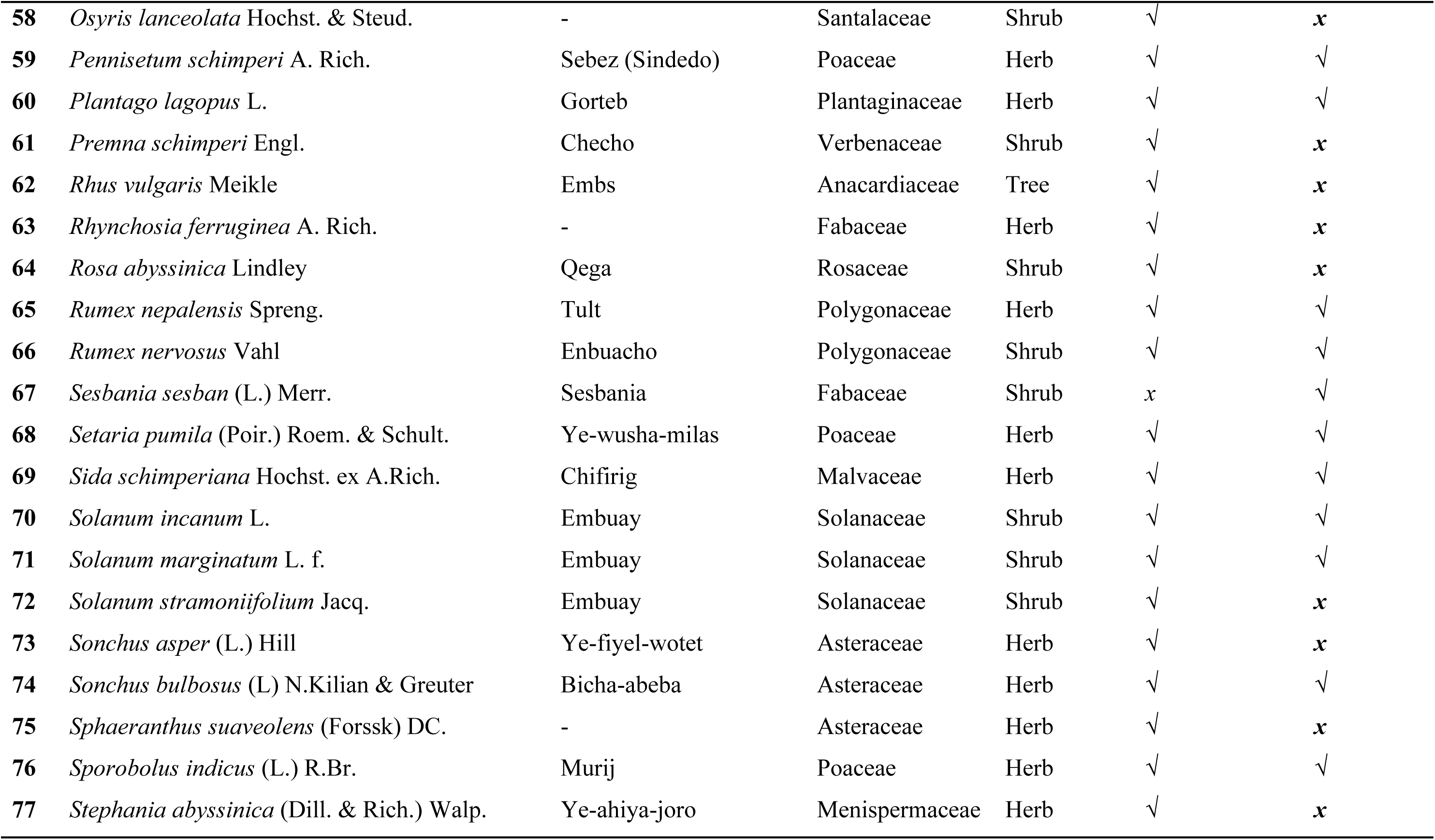
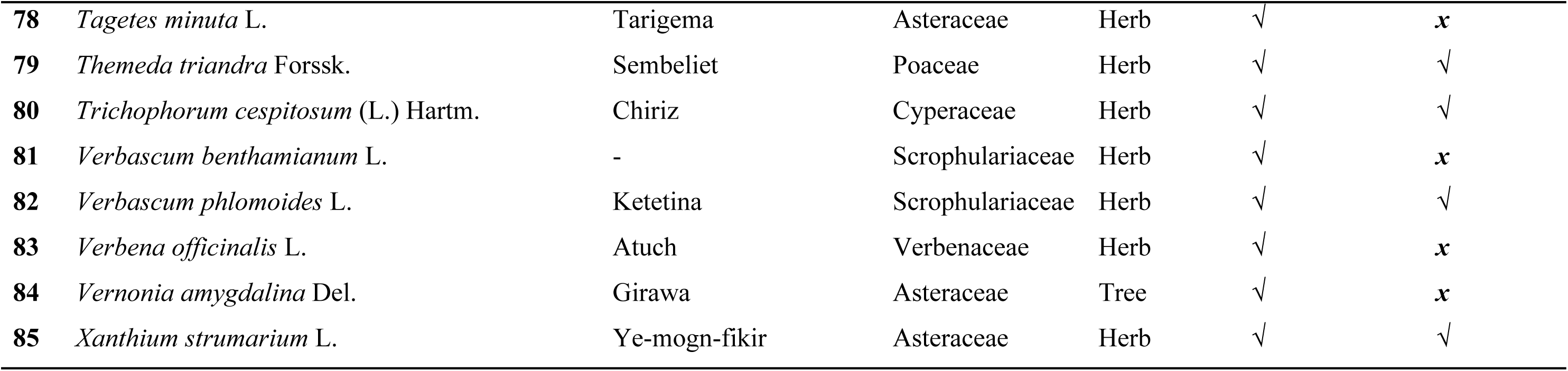

| No. | Botanical Name | Local Amharic Name | Family Name | Life Forms | Non-invaded | Invaded |
| --- | --- | --- | --- | --- | --- | --- |
| 1 | *Acacia mearnsii* De Wild. | Shiferie | Fabaceae | Tree | √ | **x** |
| 2 | *Acacia tortilis* (Forssk.) Hayne | Girar | Fabaceae | Tree | √ | √ |
| 3 | *Acanthus arboreus* Forssk. | Kosheshila | Acanthaceae | Shrub | √ | √ |
| 4 | *Achyranthes aspera* L. | Talezhe | Amaranthaceae | Herb | √ | **x** |
| 5 | *Acmella caulirhiza* Del. | Yemdir-beberie | Asteraceae | Herb | √ | **x** |
| 6 | *Agave sisalana* Perrine | Chiret (Qacha) | Asparagaceae | Shrub | √ | √ |
| 7 | *Ageratum conyzoides* (L.) L. | - | Asteraceae | Herb | √ | √ |
| 8 | *Andropogon distachyos* L. | Gaya | Poaceae | Herb | √ | √ |
| 9 | *Argemone mexicana* L. | Nech-lebash | Papaveraceae | Herb | √ | **x** |
| 10 | *Artemisia ludoviciana* Nutt. | - | Asteraceae | Herb | √ | **x** |
| 11 | *Arthraxon hispidus* (Thunb.) Makino | Qokicha | Poaceae | Herb | √ | √ |
| 12 | *Bidens pilosa* L. | Ysatan-merfy | Asteraceae | Herb | √ | √ |
| 13 | *Brucea antidysenterica* J.F.Mill. | Ye-dega-abalo | Simaroubaceae | Tree | √ | √ |
| 14 | *Caesalpinia decapetala* (Roth) Alston | Qontir | Fabaceae | Shrub | √ | √ |
| 15 | *Calpurina aurea* (Ait.) Benth. | Digita | Fabaceae | Shrub | √ | **x** |
| 16 | *Carissa spinarum* L. | Agam | Apocynaceae | Shrub | √ | √ |
| 17 | *Carthamus lanatus* L. | Ye-ahiya-suf | Asteraceae | Herb | √ | √ |
| 18 | *Cirsium vulgare* (Savi) Ten. | Ye-ahiya-shoh | Asteraceae | Herb | √ | **x** |
| 19 | *Clematis hirsuta* Perr. & Guill. | Tero | Ranunculaceae | Herb | √ | **x** |
| 20 | *Cordia africana* Lam. | Wanza | Boraginaceae | Tree | √ | √ |
| 21 | *Crepis dioscoridis* L. | - | Asteraceae | Herb | √ | **x** |
| 22 | *Croton macrostachyus* Del. | Bisana | Euphorbiaceae | Tree | √ | √ |
| 23 | *Cucumis ficifolius* A.Rich | Yemdir-embuay | Cucurbitaceae | Herb | √ | **x** |
| 24 | *Cupressus lusitanica* Mill. | Ye-ferenji-tsid | Cupressaceae | Tree | *x* | √ |
| 25 | *Cynodon dactylon* (L.) Pers. | Serdo | Poaceae | Herb | √ | √ |
| 26 | *Cyperus longus* L. | Engicha | Cyperaceae | Herb | √ | √ |
| 27 | *Desmodium intortum* (Mill.) Urb. | - | Fabaceae | Herb | √ | **x** |
| 28 | *Digitaria sanguinalis* (L.) Scop. | Warat | Poaceae | Herb | √ | √ |
| 29 | *Dodonaea viscosa* (L.) Jacq. | Kitkita | Sapindaceae | Shrub | √ | **x** |
| 30 | *Echinops kebericho* Mesfin | Kebericho | Asteraceae | Herb | √ | √ |
| 31 | *Echinops macrochaetus* Fresen | - | Asteraceae | Herb | √ | √ |
| 32 | *Echium plantagineum* L. | Aradaw | Boraginaceae | Herb | *x* | √ |
| 33 | *Eleusine floccifolia* (Forssk.) Spreng. | Akrima | Poaceae | Herb | √ | √ |
| 34 | *Erigeron bonariensis* L. | - | Asteraceae | Herb | √ | **x** |
| 35 | *Eucalyptus globulus* Labill. | Nech-bahirzaf | Myrtaceae | Tree | √ | √ |
| 36 | *Euphorbia candelabrum* Trema. | Qulqual | Euphorbiaceae | Tree | *x* | √ |
| 37 | *Ficus palmata* Forssk. | Beles | Moraceae | Shrub | √ | **x** |
| 38 | *Ficus vasta* Forssk. | Warka | Moraceae | Tree | √ | √ |
| 39 | *Foeniculum vulgare* Mill. | Ensilal | Apiaceae | Shrub | √ | √ |
| 40 | *Galinsoga parviflora* Cav. | Dha-neqay | Asteraceae | Herb | √ | √ |
| 41 | *Guizotia scabra* (Vis.) Chiov | Mech | Asteraceae | Herb | √ | √ |
| 42 | *Hackelia virginiana* (L.) I.M. Johnst. | Shingug | Boraginaceae | Herb | √ | √ |
| 43 | *Haplocoelum foliolosum* (Hiem) Bullock | Adey-ababa | Sapindaceae | Shrub | √ | √ |
| 44 | *Hygrophila schulli*. (Hamilt.) M.R. Almeida & S.M. Almeida | Amiekala | Acanthaceae | Herb | √ | √ |
| 45 | *Hyparrhenia hirta* (L.) Stapf | Sembeliet | Poaceae | Herb | √ | √ |
| 46 | *Justicia schimperiana* (Hochst. ex Nees) | Sensel | Acanthaceae | Shrub | √ | √ |
| 47 | *Lactuca serriola* L. | Enzhimo | Asteraceae | Herb | √ | √ |
| 48 | *Laggera crispata* (Vahl) Hepper & J.R.I.Wood. | - | Asteraceae | Herb | √ | **x** |
| 49 | *Laphangium luteoalbum* (L.) Tzvelev | Ye-ahiya-arti | Asteraceae | Herb | √ | √ |
| 50 | *Leucas martinicensis* (Jacq.) R.Br. | Ras-kimir | Lamiaceae | Herb | √ | **x** |
| 51 | *Lippia adoensis* Hochst. ex Walp. | Kessie | Verbenaceae | Shrub | √ | **x** |
| 52 | *Maytenus arbutifolia*. (A. Rich) Wilezek. | Atat | Celastraceae | Shrub | **x** | √ |
| 53 | *Medicago polymorpha* L. | Maget | Fabaceae | Herb | √ | √ |
| 54 | *Melinis repens* (Willd.) Zizka | - | Poaceae | Herb | √ | √ |
| 55 | *Mentha rotundifolia* (L.) Huds. | Gime | Lamiaceae | Herb | √ | √ |
| 56 | *Nuxia congesta* Fresen. | Keskeso | Loganiaceae | Shrub | √ | **x** |
| 57 | *Orthosiphon schimperi* Benth. | - | Lamiaceae | Herb | √ | √ |
| 58 | *Osyris lanceolata* Hochst. & Steud. | - | Santalaceae | Shrub | √ | **x** |
| 59 | *Pennisetum schimperi* A. Rich. | Sebez (Sindedo) | Poaceae | Herb | √ | √ |
| 60 | *Plantago lagopus* L. | Gorteb | Plantaginaceae | Herb | √ | √ |
| 61 | *Premna schimperi* Engl. | Checho | Verbenaceae | Shrub | √ | **x** |
| 62 | *Rhus vulgaris* Meikle | Embs | Anacardiaceae | Tree | √ | **x** |
| 63 | *Rhynchosia ferruginea* A. Rich. | - | Fabaceae | Herb | √ | **x** |
| 64 | *Rosa abyssinica* Lindley | Qega | Rosaceae | Shrub | √ | **x** |
| 65 | *Rumex nepalensis* Spreng. | Tult | Polygonaceae | Herb | √ | √ |
| 66 | *Rumex nervosus* Vahl | Enbuacho | Polygonaceae | Shrub | √ | √ |
| 67 | *Sesbania sesban* (L.) Merr. | Sesbania | Fabaceae | Shrub | x | √ |
| 68 | *Setaria pumila* (Poir.) Roem. & Schult. | Ye-wusha-milas | Poaceae | Herb | √ | √ |
| 69 | *Sida schimperiana* Hochst. ex A.Rich. | Chifirig | Malvaceae | Herb | √ | √ |
| 70 | *Solanum incanum* L. | Embuay | Solanaceae | Shrub | √ | √ |
| 71 | *Solanum marginatum* L. f. | Embuay | Solanaceae | Shrub | √ | √ |
| 72 | *Solanum stramonifolium* Jacq. | Embuay | Solanaceae | Shrub | √ | **x** |
| 73 | *Sonchus asper* (L.) Hill | Ye-fiyel-wotet | Asteraceae | Herb | √ | **x** |
| 74 | *Sonchus bulbosus* (L) N.Kilian & Greuter | Bicha-abeba | Asteraceae | Herb | √ | √ |
| 75 | *Sphaeranthus suaveolens* (Forssk) DC. | - | Asteraceae | Herb | √ | **x** |
| 76 | *Sporobolus indicus* (L.) R.Br. | Murij | Poaceae | Herb | √ | √ |
| 77 | *Stephania abyssinica* (Dill. & Rich.) Walp. | Ye-ahiya-joro | Menispermaceae | Herb | √ | **x** |
| **78** | *Tagetes minuta* L. | Tarigema | Asteraceae | Herb | √ | **x** |
| **79** | *Themeda triandra* Forssk. | Sembeliet | Poaceae | Herb | √ | √ |
| **80** | *Trichophorum cespitosum* (L.) Hartm. | Chiriz | Cyperaceae | Herb | √ | √ |
| **81** | *Verbascum benthamianum* L. | - | Scrophulariaceae | Herb | √ | **x** |
| **82** | *Verbascum phlomoides* L. | Ketetina | Scrophulariaceae | Herb | √ | √ |
| **83** | *Verbena officinalis* L. | Atuch | Verbenaceae | Herb | √ | **x** |
| **84** | *Vernonia amygdalina* Del. | Girawa | Asteraceae | Tree | √ | **x** |
| **85** | *Xanthium strumarium* L. | Ye-mogn-fikir | Asteraceae | Herb | √ | √ |

## Notes

### Competing Interest Statement

The authors have declared no competing interest.

